# Lempel-Ziv complexity of simultaneous surface electromyography and magnetomyography during muscle fatigue

**DOI:** 10.64898/2026.03.11.711052

**Authors:** Lorenzo Semeia, Haodi Yang, Manuela Zimmer, Markus Siegel, Oliver Röhrle, Justus Marquetand

## Abstract

**Objective:** Complexity-based metrics have been applied to surface electromyography (sEMG) to characterize fatigue-related changes in the temporal structure of myoelectric signals beyond amplitude and spectral features. Optically pumped magnetometers (OPM) are sensors that enable non-invasive recordings of magnetomyographic (MMG) signals from skeletal muscle and are increasingly used to complement surface electromyography; however, it remains unclear whether complexity measures derived from magnetic recordings are comparable to those obtained from sEMG. Here, we directly compared fatigue-related dynamics of conventional and complexity-based signal features of sEMG and OPM-MMG measured from the biceps brachii during sustained elbow flexion.

**Approach:** Healthy participants performed isometric contractions at 20% maximal voluntary contraction (MVC; 20 min) or 60% MVC (3 min). sEMG and OPM-based MMG were recorded simultaneously, and signal median frequency, root mean square (RMS), and Lempel-Ziv (LZ) complexity were calculated over time.

**Main results:** Across contraction intensities, sEMG and MMG showed consistent fatigue-related changes, characterized by increasing RMS, decreasing median frequency, and a progressive decline in LZ over time. In addition, multiple regression analyses indicated that the decrease in LZ was not fully accounted for by concurrent amplitude or spectral changes, suggesting that complexity captures aspects of signal organization that are not fully explained by established features. Finally, while sEMG showed higher LZ complexity and median frequency at 60% compared to 20% MVC, corresponding intensity-dependent effects were not observed in OPM-based MMG.

**Significance:** These findings suggest that complexity-based metrics capture fatigue-related changes in neuromuscular signal organization beyond conventional measures, and that sEMG and OPM-based MMG provide similar, though modality-specific, information. Together, the results support the use of complexity metrics in multimodal electrophysiological and biomagnetic assessments of neuromuscular fatigue.

## 1. Introduction

Human motor output emerges from interacting processes that span multiple spatial and temporal scales, from motor unit recruitment and synchronization to the biomechanics of force transmission [1]. As a result, muscle activity is typically nonlinear and exhibit structured variability rather than simple noise. This has motivated the use of complexity-based metrics to characterize myoelectric activity [2]. “Complexity” refers to structured dynamics that emerge from interactions across multiple components and scales, and are not fully captured by analyses of individual components in isolation. Different indicators of signal complexity, usually quantified using entropy- or compressibility-based measures, have been applied to surface electromyography (sEMG) to characterize changes in the temporal organization of myoelectric activity that are not fully captured by conventional amplitude or spectral features [3–7]. As such, complexity-based analyses of sEMG have been applied to aging [8] and neurological disorders such as Parkinson’s disease [9] or stroke [10], indicating sensitivity to clinically relevant alterations in motor output.

In addition to these application areas, sustained muscle contractions provide a well-controlled experimental setup for examining how the temporal organization of muscle activity changes over the course of sustained activation and the development of fatigue. In neuromuscular physiology, muscle fatigue is a multifactorial process arising from interactions between central and peripheral mechanisms that alter motor unit activation, force production, and neuromuscular control during sustained or repeated contractions [11]. Classical electrophysiological fatigue markers derived from sEMG include, for example, amplitude-based measures reflecting overall signal amplitude, which typically increase over time due to compensatory motor unit recruitment and firing rate adjustments [12]. Moreover, spectral features typically shift toward lower frequencies during sustained isometric contractions and are linked to fatigue-related changes in muscle fiber conduction velocity [13]. While these fatigue-related markers are consistently observed across muscle groups, the rate of their evolution over time depends on contraction intensity, with steeper fatigue-related slopes at higher force levels [14].

Beyond these parameters, fatigue also alters the temporal organization of electromyographic signals. This organization is summarized using complexity metrics, which are commonly operationalized through entropy- or compressibility-based measures. Such measures have been applied to sEMG to capture fatigue-related changes not fully explained by amplitude or spectral features [2]. In this context, sustained contractions are typically associated with a progressive reduction in complexity over time [15]. In addition, sEMG complexity increases from low to moderate contraction intensities but shows little further increase at higher force levels, consistent with a plateauing effect that may arise when force production is predominantly regulated by motor unit discharge rate [15]. Among the different approaches, Lempel-Ziv (LZ) complexity has been proposed as a computationally efficient measure to capture fatigue-related changes, as it quantifies the degree of regularity and repetition in the signal and is sensitive to alterations in myoelectric patterns not captured by conventional spectral indices [3,16]. Therefore, a decrease in LZ complexity over time indicates an increasingly regular myoelectric pattern, reflecting less variable neuromuscular output as fatigue develops.

While sEMG reflects voltage differences at the skin surface, muscle activation also generates biomagnetic fields that can be measured as magnetomyographic signals. Magnetic recordings of skeletal muscle activity have long been discussed as a valuable physiological measurement, with sensitivity to aspects of action potential propagation and spatial current distributions that are not directly accessible with sEMG [17]. Technical development in optically pumped magnetometers (OPM)-based magnetomyography (MMG) has substantially increased the feasibility of muscle biomagnetism by enabling room-temperature, wearable sensors that can be positioned close to the muscle [18] and measured in portable magnetic shields [19]. Moreover, OPM-based MMG has been shown to provide complementary information to sEMG rather than simply being redundant with it [20,21]. For example, while both sEMG and MMG have been shown to exhibit a shift of the signal spectrum toward lower frequencies during fatigue, magnetic recordings provide additional information by enabling access to spatial and directional aspects of muscle activity that are not captured by surface electromyography [22].

However, while sEMG and MMG show comparable behavior for conventional frequency- and amplitude-based features, it remains unclear whether these similarities extend to metrics quantifying signal complexity. As OPM-based MMG is increasingly considered a valuable tool for neuromuscular research in healthy and pathological conditions [23,24], it is important to assess not only established spectral markers but also the richness and temporal organization of the recorded signals, which may capture complementary aspects of neuromuscular dynamics. Here, we examine whether complexity-based and conventional fatigue markers show coherent temporal dynamics across sEMG and MMG recordings obtained during sustained isometric elbow flexion contractions. We focus on three complementary descriptors computed in short time windows: LZ complexity as an index of temporal structure, median frequency as a spectral fatigue marker, and root mean square (RMS) as an amplitude-related measure of muscle activation. Two contraction intensities (20% MVC and 60% MVC) were analyzed to assess intensity-dependent effects. Based on this approach, we pursued three aims: first, to characterize how neuromuscular signal complexity evolves over time during fatigue in both sEMG and OPM-based MMG; second, to assess whether complexity captures aspects of signal organization that are not fully described by conventional spectral or amplitude-based indices; and third, to examine how these relationships depend on contraction intensity. Based on previous works, we expected that LZ complexity would decrease over time during sustained contractions, that the rate of change would be steeper at higher contraction intensities, and that comparable temporal patterns would be observed in both sEMG and OPM-based MMG.

## 2. Methods

### 2.1. Participants and recordings

Two experimental protocols in healthy young adults were included in this manuscript, both involving isometric elbow flexion. In the first condition, *n* = 12 participants (6 females) performed a sustained contraction at 20% of their maximal voluntary contraction (MVC) for 20 minutes. In the second condition, *n* = 6 participants (3 females) completed a 3-minute contraction at 60% MVC. The study was conducted in accordance with the Declaration of Helsinki and approved by the local ethics committee; written informed consent was obtained from all participants.

### 2.2. Experimental setup and data acquisition

All recordings were acquired at the MEG Center of the University of Tübingen, inside a magnetically shielded room (Ak3b, VAC Vacuumschmelze, Hanau, Germany). Neuromuscular activity was recorded from the biceps brachii muscle using sEMG and OPM-based MMG. One channel sEMG (Conmed, Cleartrace2 MR-ECG-electrodes) was measured using a bipolar configuration. The ground and the reference electrodes were placed on the right shoulder and on the ipsilateral proximal forearm. A single OPM sensor (QZFM-gen-1.5, QuSpin Inc., Louisville, CO, USA) was used, and one magnetic field component was included in the analysis. The analog outputs of the OPM sensor were measured using the electronics of a CTF Omega 275 MEG system (Coquitlam, BC, Canada). The system’s EEG input channels were simultaneously used to record the sEMG signals. Both sEMG and OPM data were sampled at *fs* = 2343.8 Hz. Prior to each measurement, ultrasound imaging (Mindray TE7, 14Mhz-linear probe) was performed by a trained physician (JM) to precisely localize the biceps muscle belly. This anatomical landmark was marked on the skin and used to guide the placement of sEMG electrodes and OPM sensor. Participants were seated comfortably with their dominant arm supported in a semi-flexed position and grabbing an in-house-built force transducer designed to measure isometric elbow-flexion force. The force signal was digitized together with sEMG and OPM recordings. At the beginning of each session, participants performed three MVCs of elbow flexion, each lasting approximately 5 seconds and separated by one minute of rest. The mean force across the three trials was used as the individual MVC reference value to define the target force levels for the subsequent fatigue experiments. In the fatigue protocol, after a brief ramp-up phase, participants were instructed to maintain a constant isometric contraction at either 20% MVC for 20 minutes or 60% MVC for 3 minutes. The study protocol is illustrated in Fig. 1.

**Fig. 1.**
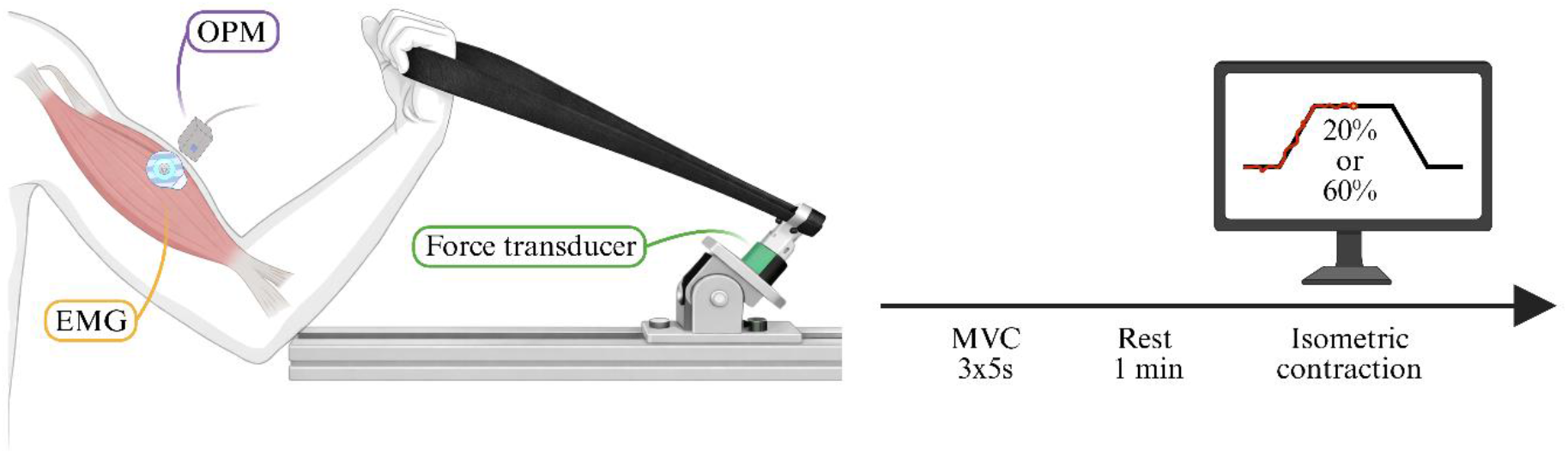
Experimental protocol. Neuromuscular activity of the biceps brachii during elbow flexion was recorded simultaneously using surface electromyography (sEMG) and an optically pumped magnetometer (OPM) placed over the muscle belly. Participants performed isometric elbow flexion while seated, with the forearm stabilized and force measured using a transducer. Maximum voluntary contraction (MVC) was recorded in three trials and the average was used to set the target force levels. During the fatigue protocol, participants maintained contractions at either 20% MVC for 20 min or 60% MVC for 3 min. Created with BioRender.com.

### 2.3. Preprocessing

All analyses were performed in MATLAB R2023b (The MathWorks, Natick, MA, USA) using FieldTrip [25]. The same pipeline was applied to both sEMG and OPM signals. Raw datasets were initially demeaned to remove DC offsets introduced by sensors and acquisition hardware. Signals were high-pass filtered at 30 Hz and low-pass filtered at 200 Hz using zero-phase, 4th-order IIR Butterworth filters. Powerline contamination and its harmonics were attenuated using 4th-order IIR Butterworth band-stop filter (frequencies: 49-51, 99-101, 149-151, 199-201 Hz). At this point, filtered signals were visually inspected, and artifact-contaminated segments were manually annotated using interactive marking and later removed. Following artifact rejection, signals were z-scored, trimmed to the isometric-contraction period, and resampled to 1000 Hz using an integer-ratio resampling scheme (125/293 × *fs* ≈ 1000 Hz).

### 2.4. Parameters estimation

To estimate temporal changes in neuromuscular activity during sustained contraction, each preprocessed time series was divided into non-overlapping 5-second windows, each consisting of 5000 time points. Windows containing removed artifacts were skipped. For each window, we computed the following parameters: Lempel-Ziv (LZ) compressibility [26], median spectral frequency, and the signal root mean square (RMS). We quantified LZ complexity using the original compression-based formulation applied to a binarized version of the sEMG and OPM signals [26]. The basic principle of the method is to scan the binary sequence and count the number of new, non-repeating substrings that appear as the sequence unfolds, with a greater count indicating higher signal complexity. For binarization, each segment was thresholded at its median value. The resulting LZ value was then normalized by *T*/log_2_(*T*), where *T* denotes the window length, to obtain an estimate of the signal’s entropy rate [27]. Power spectral densities were obtained using Welch’s method (700-sample windows). The cumulative spectral distribution was used to determine the median frequency. Finally, the RMS of each 5-second segment was calculated to capture amplitude-related fatigue effects.

For each participant and each parameter, a quadratic polynomial was fit to the window-wise trajectory. Residuals exceeding three standard deviations were labelled as outliers and replaced with *NaN*. Cleaned parameter time series were then used for all subsequent analyses.

### 2.5. Statistical analysis

Our first aim was to characterize the pairwise relationships among neuromuscular parameters over time by computing correlation matrices across all sEMG- and OPM-based MMG-derived measures (LZ, median frequency, and RMS). Two correlation matrices were computed, one for the 20% MVC protocol and one for the 60% MVC protocol. These matrices allowed us to assess the extent to which different features correlated during muscle activity and to quantify shared variances across modalities. For this analysis, window-wise parameter trajectories were first averaged across participants to obtain group-level time courses for each metric. Pearson correlation coefficients were then calculated using MATLAB’s “corrcoef*”* function (pairwise deletion), with statistical significance assessed at an alpha level of 0.05.

Next, to isolate the unique contribution of each neuromuscular parameter, we modeled each feature within each modality (sEMG or OPM-MMG) using a set of linear models. For these analyses, the dependent variable (LZ, median frequency, or RMS) was regressed onto time and the remaining two parameters within the same modality. All measures were z-scored prior to modeling in order to place predictors on a comparable scale and allow the regression coefficients to be interpreted as standardized effects. This approach enabled us to disentangle direct time-related fatigue effects from cross-parameter dependencies. Regression coefficients, 95% confidence intervals, and p-values were extracted from each model.

Finally, to compare fatigue dynamics across contraction intensities, we pooled data from the 20% and 60% MVC protocols and fit a series of linear mixed-effects models. For each parameter and modality, we modeled the window-wise value as a function of protocol (20% *vs*. 60% MVC), normalized time (0-1), and their interaction. This allowed us to determine whether overall parameter levels differed between protocols, whether fatigue progressed over time, and whether the rate of fatigue-related change differed between intensities. Predicted trajectories with 95% confidence intervals were used to visualize protocol differences.

To correct for multiple testing, we applied the false discovery rate (FDR) correction procedure by Benjamini and Hochberg [28]. This correction was performed separately for each family of analyses, namely: (1) the correlation matrix for the 20% MVC protocol, (2) the correlation matrix for the 60% MVC protocol, (3) the linear models results for the 20% MVC condition, (4) the linear models results for the 60% MVC condition, and (5) the fixed-effect terms from the mixed-effects models comparing the two contraction intensities.

## 3. Results

### 3.1. Dataset quality and preprocessing

We initially included 12 participants in the 20% MVC protocol and 6 participants in the 60% MVC protocol. In the 20% MVC condition, one dataset was excluded due to detachment of an sEMG electrode, leaving 11 usable datasets. In the 60% MVC condition, one participant was unable to maintain the target force level and was therefore excluded, resulting in 5 usable datasets. In addition, one participant in the 60% MVC group displayed abnormally low RMS in the sEMG and was excluded specifically from analyses involving RMS. After removing the ramp-up and ramp-down periods, the duration of the sustained contraction segments was either 3 or 20 minutes across participants, yielding up to 36 or 240 non-overlapping clean 5-second windows per subject.

We excluded short artifact-contaminated segments identified during visual inspection (sEMG: 6 segments across 2 recordings, median duration 5.8 seconds; OPM: 10 segments across 8 recordings, median duration 6.2 seconds). Following artifact removal, a quadratic polynomial was fitted to each parameter trajectory over time, and values whose residuals exceeded 3 standard deviations were classified as outliers. In the 20% MVC condition, 3.3% of all data points were removed as outliers across modalities and parameters; in the 60% MVC condition, 0.9% were removed.

### 3.2. Correlation analyses

To characterize dynamics across neuromuscular signals, we computed pairwise Pearson correlation matrices including time and all sEMG- and OPM-derived parameters (LZ complexity, median frequency, and RMS), separately for the 20% MVC (Fig. 2A) and 60% MVC (Fig. 2B) contractions. Parameters were first computed for each subject and time window and then averaged across participants to obtain group-level time courses, from which the correlation matrices were derived. Correlations with time quantified the direction and strength of temporal changes in each parameter, while correlations among parameters captured their shared temporal structure. In addition, the matrices captured both within-modality (sEMG-sEMG, OPM-OPM) and cross-modality (sEMG-OPM) relationships, allowing us to assess how consistently the two sensor types tracked fatigue-related changes over time. All Pearson’s r values and FDR-corrected significance are shown in the correlation matrices. At both 20% and 60% MVC, LZ complexity and median frequency declined over time, whereas RMS increased, reflecting the classical spectral slowing and amplitude augmentation expected during sustained isometric fatigue (Fig. 3). Importantly, these temporal patterns were observed in both sEMG and OPM measurements, indicating that the two sensor types capture highly similar physiological processes during fatigue. As shown in the correlation matrices, LZ complexity was also strongly related to both median frequency and RMS, consistent with the idea that complexity reductions emerge alongside spectral compression and amplitude increases. However, correlation analyses do not distinguish whether variations in LZ are associated primarily with changes in spectral content, signal amplitude, or variance shared between these features. We therefore applied multiple linear regression models to isolate the unique contributions of median frequency and RMS to LZ values. This approach allowed us to assess whether LZ changes could be explained by amplitude or spectral features, or whether they exhibited an additional time-dependent component beyond their shared correlation structure.

**Fig. 2.**
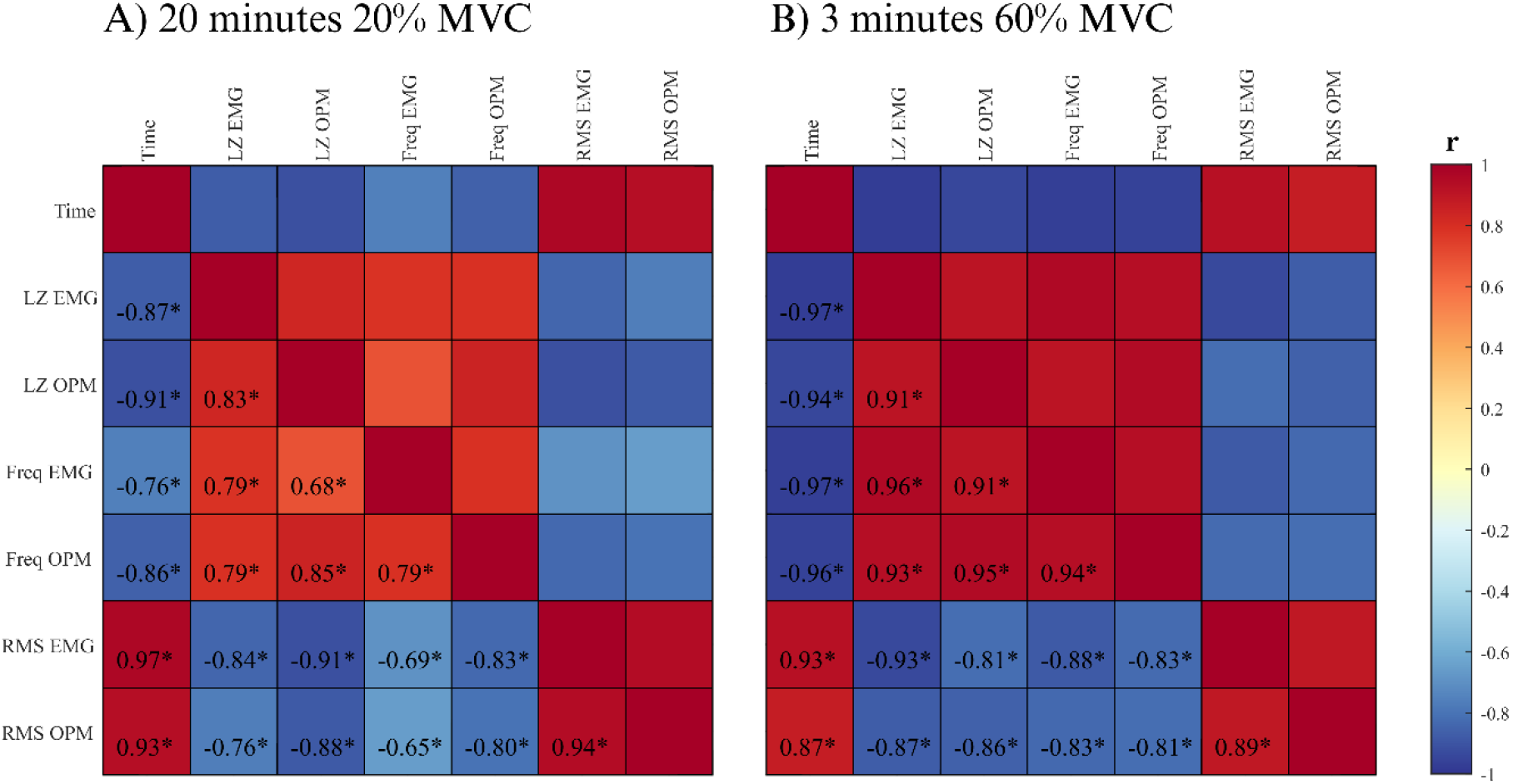
Correlation matrices of fatigue-related neuromuscular parameters. Pairwise Pearson correlation matrices showing relationships between Lempel-Ziv (LZ) complexity, median frequency, and root mean square (RMS) derived from surface electromyography (sEMG) and optically pumped magnetometers (OPM). Correlations are shown separately for the sustained 20% MVC contraction (A, 20 minutes) and the 60% MVC contraction (B, 3 minutes). Numerical values indicate Pearson’s correlation coefficients (r); asterisks denote statistically significant correlations after false discovery rate (FDR) correction (p_FDR_ < 0.05). Created with BioRender.com.

**Fig. 3.**
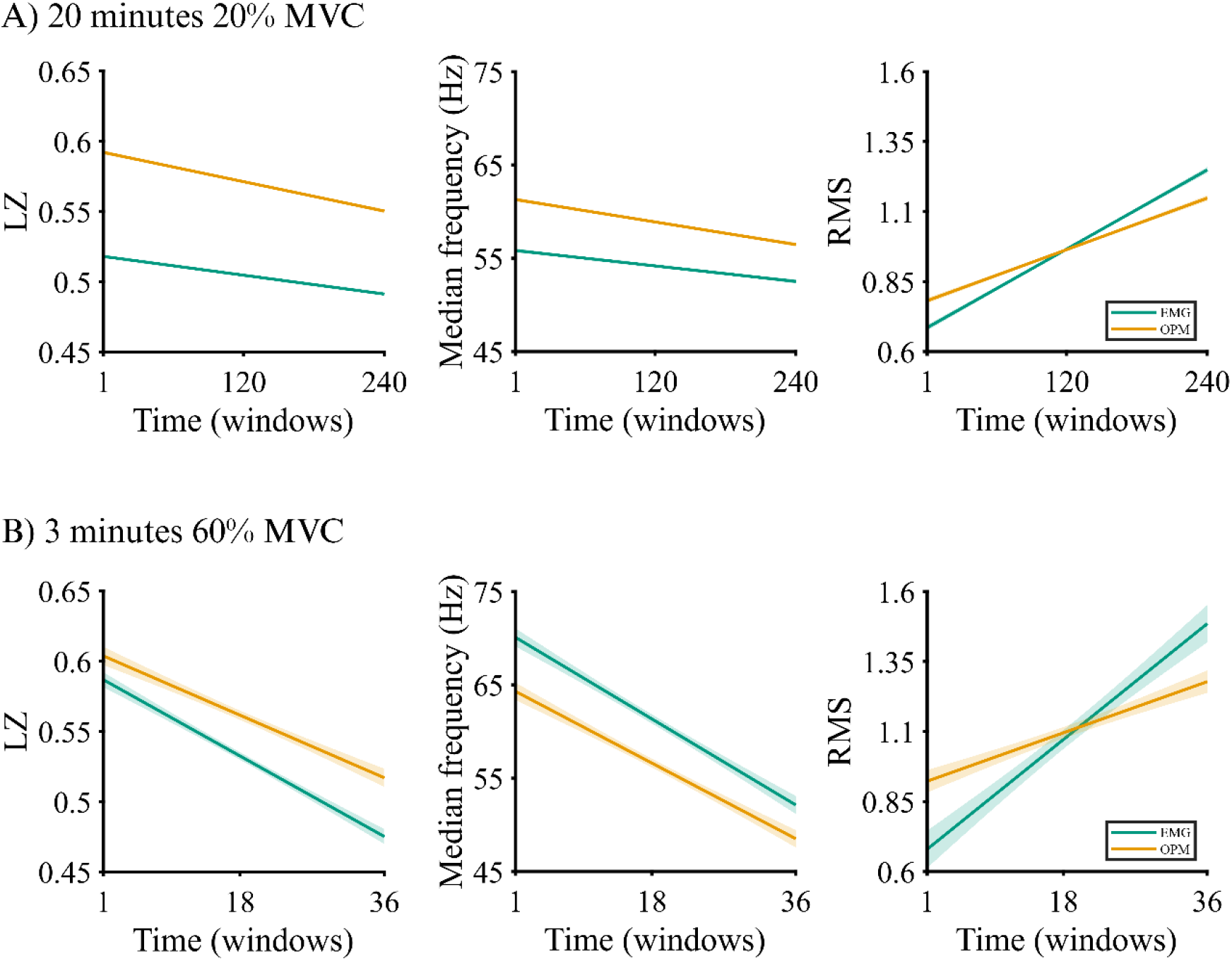
Temporal evolution of neuromuscular parameters during sustained contractions. Time courses of Lempel-Ziv (LZ) complexity, median frequency, and RMS derived from sEMG and OPM-based MMG during isometric contractions at 20% MVC (A) and 60% MVC (B). Solid lines indicate linear model fits with shaded areas representing 95% confidence intervals. All correlations are statistically significant after false discovery rate (FDR) correction (p_FDR_ < 0.05). Created with BioRender.com.

### 3.3. Partial contributions of RMS, frequency, and time to LZ complexity

To characterize how different neuromuscular features evolve over time, we fitted linear models within each modality (sEMG and OPM separately) in which each parameter (LZ, median frequency, RMS) was predicted from time and the remaining two parameters. All variables were z-scored before fitting the models so that they shared a common scale and the resulting coefficients reflected standardized effects. The analysis was conducted separately for the 20% MVC (Fig. 4A) and 60% MVC (Fig. 4B) recordings. While models were estimated for all three dependent variables, our primary interest lay in the predictors of LZ complexity.

**Fig. 4.**
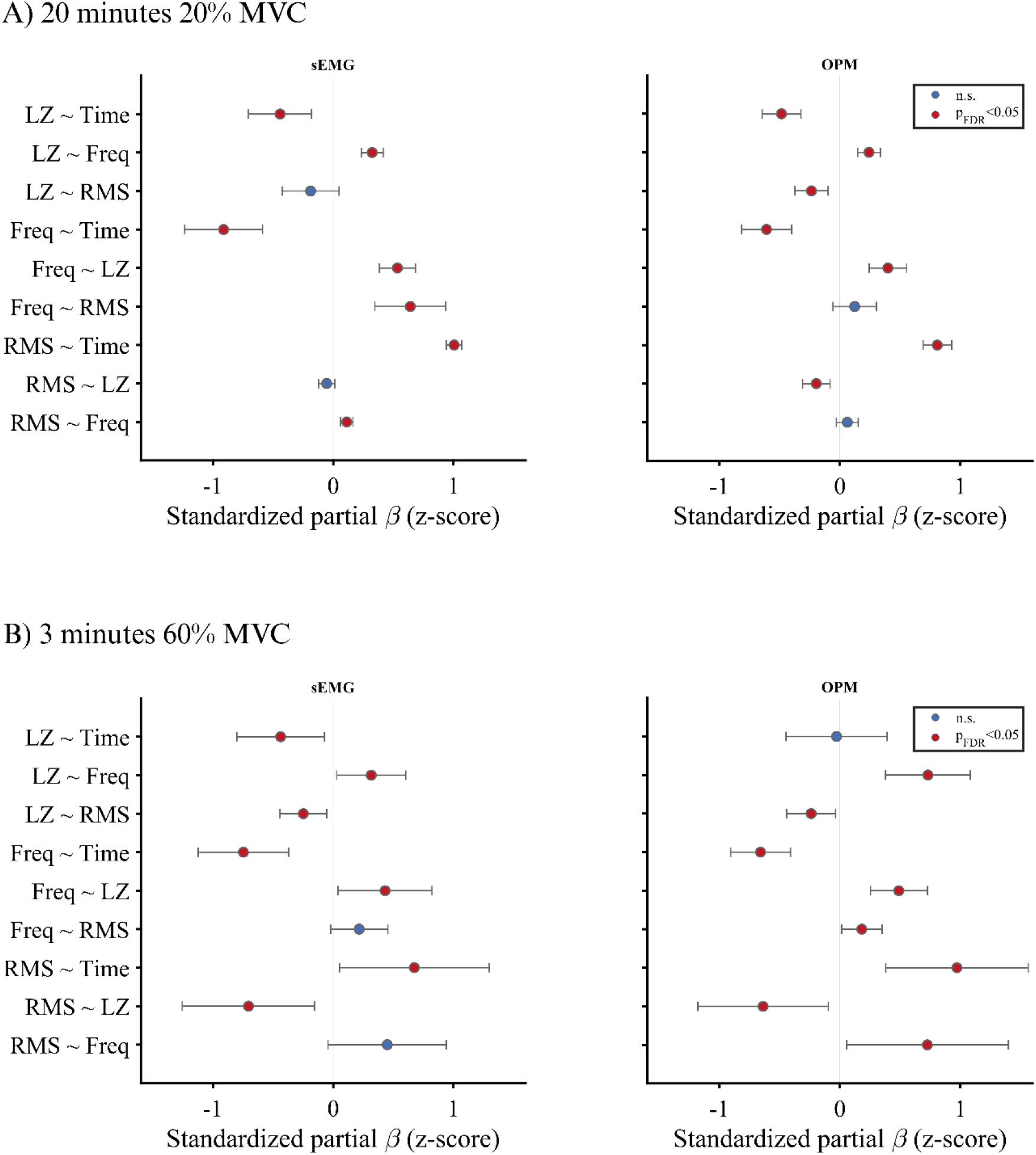
Partial contributions of time, median frequency (Freq), and RMS to fatigue-related neuromuscular parameters. Standardized partial regression coefficients (β, z-scored) from multiple linear regression models fitted separately for surface electromyography (EMG, left) and OPM-based magnetomyography (OPM, right). Models were estimated for (A) the 20% MVC, 20-min contraction and (B) the 60% MVC, 3-min contraction. Each row shows the partial association between a dependent variable (LZ complexity, median frequency, or RMS) and a given predictor (time, median frequency, or RMS), after accounting for the remaining predictors in the model. Points indicate standardized β estimates and horizontal bars denote 95% confidence intervals. Red markers indicate effects surviving false discovery rate (FDR) correction (p_FDR_<0.05); blue markers indicate non-significant effects. Created with BioRender.com.

In the 20% MVC condition, LZ showed positive partial associations with median frequency (β = 0.32, p_FDR_ < 0.001) and negative partial associations with RMS, although not statistically significant (β = - 0.19, p_FDR_ = 0.13), indicating that complexity was generally lowest when the signal became stronger in amplitude and spectrally slower, which are typical markers of fatigue. Importantly, time remained a significant predictor of LZ even after adjusting for median frequency and RMS (β = -0.44, p_FDR_ = 0.001), suggesting that the progressive decline in complexity cannot be fully accounted for by spectral changes or amplitude alone. Instead, LZ appears to capture an additional, time-dependent aspect of signal structure that is not related to classical sEMG features. A similar pattern was observed for the OPM recordings, in which LZ also showed a positive partial association with median frequency (β = 0.24, p_FDR_ < 0.001) and negative partial associations with RMS (β = -0.23, p_FDR_ = 0.001) and time (β = -0.48, p_FDR_< 0.001). These results support the idea that OPM measurements can capture fatigue-related effects similarly to what is observed in the sEMG signal. Although median frequency and RMS were also modeled as dependent variables, their effects primarily reflected well-established fatigue trends; for completeness, these numerical results (including regression coefficients, 95% confidence intervals, and p_FDR_ values) are reported in Sup. Table 1. Standardized partial regression coefficients are also visualized as forest plots in Fig. 4A.

Finally, the same analysis was conducted in 60% MVC recordings, and it yielded a qualitatively similar pattern of associations (Fig. 4B, Sup. Table 2).

### 3.4. Comparison between the 20% and 60% MVC contractions

To directly compare fatigue-related trajectories across contraction intensities, we fitted linear mixed-effects models to the sEMG and OPM-derived parameters (LZ, median frequency, RMS), using the respective protocols (20% *vs*. 60% MVC), normalized time (0-1), and their interaction as predictors. In the sEMG signals, for LZ complexity (Fig. 5A), both protocol (β = 0.07, p_FDR_ < 0.0001) and time (β = -0.03, p_FDR_ = 0.009) contributed significantly to the model, and a significant protocol × time interaction (β = -0.08, p_FDR_ < 0.0001) indicates that LZ declined more steeply during the 60% *vs*. 20% MVC contraction. Median frequency (Sup. Fig. 1A) showed a similar pattern, with a main effect of protocol (β = 14.11, p_FDR_ <0.0001) and a significant interaction protocol × time (β = 14.49, p_FDR_ = 0.0001), but no effect of time (p_FDR_ > 0.05) also suggesting a faster decline in frequency over time in the 60% *vs*. 20% MVC condition. RMS (Sup. Fig. 2A) showed a general increase over time (β = 0.56, p_FDR_ < 0.0001), with no significant effect of protocol (p_FDR_ > 0.05) nor protocol × time interaction (p_FDR_ > 0.05). Results for the OPM-derived signals (Fig. 5B, Sup. Fig. 1B, Sup. Fig. 2B) showed qualitatively similar patterns, with comparable directions of effect for protocol, time, and protocol × time. Full model coefficients and significance values are reported in Sup. Table 3.

**Fig. 5.**
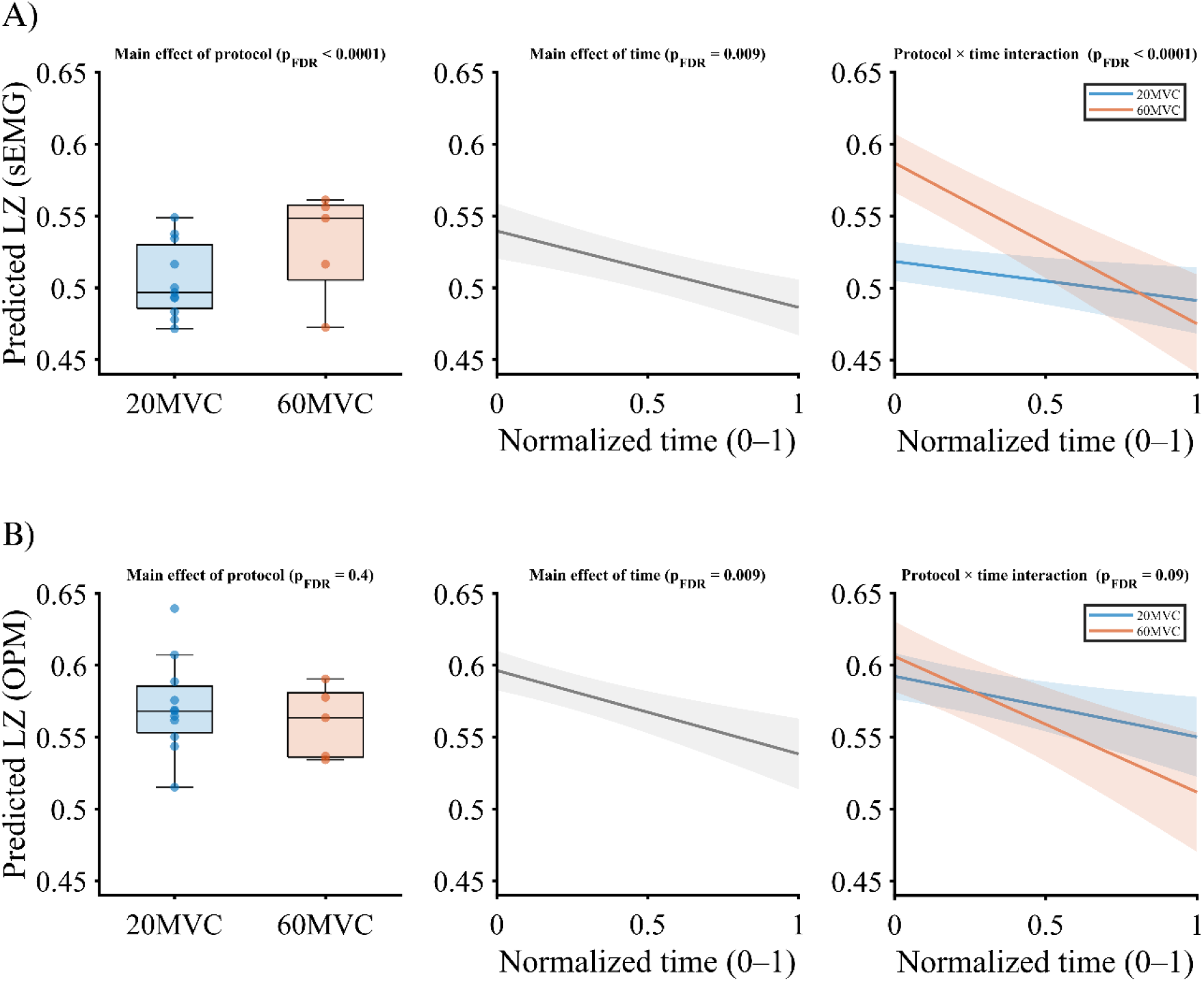
Intensity- and time-dependent effects on Lempel-Ziv (LZ) complexity in sEMG and OPM recordings. Predicted LZ values derived from linear mixed-effects models are shown for surface electromyography (sEMG; A) and OPM-based magnetomyography (OPM; B). Left panels depict the main effect of protocol (20% vs. 60% MVC), middle panels show the main effect of normalized time (0-1), and right panels illustrate the protocol × time interaction. Boxplots show model-predicted values aggregated across time. Solid lines represent model-predicted trajectories, with shaded areas indicating 95% confidence intervals. False discovery rate (FDR)-corrected p-values are reported above each panel. Created with BioRender.com.

## 4. Discussion

In this study, we investigated fatigue-related changes in neuromuscular signal complexity derived from surface electromyography (sEMG) and optically pumped magnetometer (OPM)-based magnetomyography (MMG) during sustained isometric contractions at two force levels. By combining classical amplitude- and frequency-based fatigue markers with Lempel-Ziv (LZ) complexity, we had three aims: first, to characterize how signal complexity evolves over time during fatigue in both sEMG and MMG; second, to evaluate whether complexity provides information about signal organization beyond that conveyed by conventional spectral or amplitude-based metrics; and third, to assess how these relationships are influenced by contraction intensity. Our results indicate that both sEMG and MMG show coherent fatigue-related changes in LZ, spectral content, and amplitude, with LZ complexity declining progressively over time in both modalities. Importantly, this decline could not be fully explained by changes in median frequency or signal root mean square (RMS), indicating that complexity represents an additional, time-dependent aspect of neuromuscular signal organization. Finally, sEMG showed higher overall LZ complexity during high-versus low-intensity contractions, whereas OPM-derived complexity did not differ systematically between contraction intensities. Overall, our results suggest that OPM-based MMG captures fatigue-related temporal organization of neuromuscular signals in a manner broadly comparable to sEMG, while also highlighting modality-specific differences that depend on contraction intensity. This multimodal approach provides a methodological foundation for using OPM-MMG to obtain fatigue-related complexity measures and supports future studies evaluating their feasibility and interpretability in clinical and rehabilitation contexts.

### 4.1. Fatigue-related changes over time

As an initial step, we assessed pairwise correlations between time, LZ complexity, median frequency, and RMS to characterize fatigue-related temporal relationships in sEMG and OPM-based MMG, as well as relationships among the parameters themselves (Fig. 2 A-B). Separate correlation matrices were computed for the two contraction intensities (20% and 60% MVC), each including both modalities, allowing assessment of relationships within and across recording modalities. Across contraction intensities, we observed consistent fatigue-related temporal patterns (Fig. 3), characterized by a progressive increase in signal RMS and a decrease in median frequency over time, in line with established electrophysiological markers of muscle fatigue [12,13]. In parallel, LZ complexity declined over time, reflecting increasingly regular and repetitive neuromuscular signal patterns and, consistent with prior work, serving as a computationally efficient and non-stationarity-robust marker of fatigue-related changes [3]. Strong associations were also observed between fatigue markers. For example, median frequency showed a negative correlation with RMS, also consistent with fatigue-related mechanisms in which compensatory increases in neural drive and changes in motor unit behavior are accompanied by spectral slowing [29]. These preliminary findings suggest that sEMG and OPM-based MMG capture similar fatigue-related changes, with signal spectral content, amplitude, and LZ complexity showing consistent temporal patterns in both modalities. However, correlation analyses do not allow inferences about unique or independent contributions of individual parameters; i.e., they cannot determine whether reductions in LZ complexity are primarily driven by spectral slowing, increases in RMS, or other time-dependent effects. To address this point, we next examined the partial contributions of time, median frequency, and RMS to changes in complexity.

### 4.2. Partial contributions of time, median frequency and RMS to LZ complexity

To assess the independent contributions of time, median frequency, and RMS to LZ complexity, we applied multiple linear regression models that account for shared variance among predictors. Across conditions (20% *vs*. 60% MVC) and modalities (sEMG *vs*. OPM-MMG), LZ complexity tended to show positive associations with median frequency and negative associations with RMS, indicating that higher complexity was generally linked to broader spectral content and lower signal amplitude (Fig. 4 A-B). Importantly, in several conditions, time remained a significant predictor of LZ complexity even after adjusting for amplitude and spectral features, suggesting that fatigue-related decline in complexity cannot be fully explained by conventional markers alone [2,3]. Physiologically, however, because complexity is derived from global sEMG or OPM-MMG features, its decline may reflect multiple, partially overlapping mechanisms. These include peripheral changes such as fatigue-related reductions in muscle fiber conduction velocity and increased temporal overlap of motor unit action potentials [1], as well as central adaptations in neural drive, including changes in motor unit recruitment strategies or discharge behavior [11,30]. The present data do not allow these peripheral and neural contributions to be separated. However, the ability of OPM-based MMG to detect single motor unit activity [31] suggests that future studies may be able to relate complexity-based parameters more directly to motor-unit-level parameters. Accordingly, a natural next step will be to combine complexity-based analyses with more specific physiological indices of peripheral neuromuscular function, such as conduction velocity estimates, spatially resolved sEMG and OPM-MMG recordings, and motor unit-level measures of neural drive (e.g., motor unit firing rate). Integrating such markers within a similar experimental paradigm, but with higher-density sensor coverage, would allow more direct separation of peripheral and neural contributions to neuromuscular signal complexity.

### 4.3. Intensity- and modality dependent characteristics of fatigue trajectories

Contraction intensity influenced fatigue-related parameters in a modality-specific manner. In sEMG, higher contraction intensity (60% MVC) was associated with higher overall LZ complexity (Fig. 5A) and higher spectral frequency (Sup. Fig. 1) than lower contraction intensity (20% MVC). These intensity-related effects were, however, not observed in OPM-based MMG (Fig. 5B). RMS amplitude did not differ between contraction intensities in either modality (Sup. Fig. 2). While modulation of fatigue-related parameters by contraction intensity is well documented in the sEMG literature [15], the absence of corresponding intensity-dependent effects in OPM-based MMG observed here suggests modality-specific sensitivities in how neuromuscular activity is captured. Potential explanations rely on the circumstance that magnetic recordings are known to depend on factors such as sensor orientation, sensor-to-source distance, and system timing characteristics, all of which can influence signal morphology and feature estimates [32,33]. Finally, the OPM data quality might also be limited by the narrow bandwidth of most OPM (here, 135 Hz). In this context, recent work has emphasized the need for more simulation and validation studies to assess how imperfect sensor responses affect biomagnetic measurements before OPM systems can be reliably used in clinical or translational settings [34].

### 4.4. Limitations

Our findings should be interpreted within the scope of the experimental design. The study was restricted to two contraction intensities and a single muscle, which limits the generalizability of the observed fatigue-complexity relationships. In addition, the use of single-channel sEMG and OPM-based MMG limits physiological interpretation, as global measures such as LZ complexity do not allow direct inference about specific mechanisms such as motor unit recruitment strategies, synchronization, or changes in muscle fiber conduction velocity. Furthermore, the sample size was modest, which may limit statistical power and the characterization of inter-individual variability. Finally, although sEMG and OPM-based MMG showed broadly comparable fatigue-related changes in signal complexity, the absence of intensity-dependent modulation in MMG suggests that modality-specific sensitivities may influence how fatigue-related signal organization is measured. Incorporating a wider range of contraction intensities and spatially resolved recordings could help assess how time-dependent changes in signal complexity relate to peripheral and central mechanisms.

### 4.5. Conclusions

In summary, this study shows that Lempel-Ziv complexity can describe fatigue-related temporal changes in neuromuscular signals derived from both sEMG and OPM-based MMG during sustained isometric contractions. Across contraction intensities, complexity declined progressively over time, indicating increasingly regular signal organization that was not fully captured by the amplitude and spectral measures used here. These findings support the use of complexity-based measures as complementary fatigue markers that characterize temporal aspects of neuromuscular signals beyond conventional features. In addition, while sEMG showed intensity-dependent differences in complexity, such modulation was not observed in OPM-based MMG, possibly highlighting modality-specific sensitivities and current technical limitations of magnetic recordings. Together, our results support the use of complexity-based measures to characterize fatigue-related signal organization, while underscoring the need for future studies combining broader intensity ranges, spatially resolved recordings, and motor-unit-level analyses to better distinguish between peripheral and central contributions to fatigue.

## Authors contribution

L.S. designed the study, collected the data, conceptualized and conducted the analysis, prepared the figures and drafted the manuscript. H.Y. supported the analysis and revised the manuscript. M.Z. supported data collection and revised the manuscript. M.S., O.R. and J.M. supervised the work. All authors discussed the results, reviewed the manuscript and approved its final version.

## Data availability statement

Processed data and the analysis code required to reproduce the results and figures of this study will be made available on the project’s Open Science Framework (OSF, project DOI: 10.17605/OSF.IO/FPDHN) repository upon publication.

## Acknowledgements

LS and JM were supported by the European Research Council (ERC) through the Advanced Grant “qMOTION” (Grant ID: 101055186). JM received support from the Deutsche Forschungsgemeinschaft (DFG, German Research Foundation) within the priority program SPP 2311 (Grant ID: 548605919). JM and MS received funding from the Bundesministerium für Bildung und Forschung (BMBF) as part of the Future Cluster “QSens” (Grant ID: 03ZU2110FD). Additional support for this project was provided by the German Space Agency (DLR) using funds from the Federal Ministry for Economic Affairs and Climate Action (BMWK), Grant DLR 50BM2534B. We also thank the International Max Planck Research School for the Mechanisms of Mental Function and Dysfunction (IMPRS-MMFD) and the Joachim Herz Foundation for the support to LS. We thank Jürgen Dax and Dr. Joel Frohlich for technical support and conceptual feedback.

## Ethics statement

The study was carried out in accordance with the Declaration of Helsinki and received approval from the Ethics Committee of the University of Tübingen. All participants provided written informed consent for participation and for the publication of their data.

