## Supplements for "Lempel-Ziv complexity of simultaneous surface electromyography and magnetomyography during muscle fatigue"

| Modality | Dependent | Predictor | β | CI low | CI high | $p_{FDR}$ |
|---|---|---|---|---|---|---|
| **EMG** | LZ | Time | -0,44 | -0,70 | -0,18 | 0,001 |
| | LZ | Freq EMG | 0,32 | 0,23 | 0,41 | <0.0001 |
| | LZ | RMS EMG | -0,19 | -0,43 | 0,05 | 0,13 |
| | Freq | Time | -0,91 | -1,24 | -0,59 | <0.0001 |
| | Freq | LZ EMG | 0,53 | 0,38 | 0,68 | <0.0001 |
| | Freq | RMS EMG | 0,64 | 0,35 | 0,94 | <0.0001 |
| | RMS | Time | 1,01 | 0,94 | 1,07 | <0.0001 |
| | RMS | LZ EMG | -0,05 | -0,12 | 0,01 | 0,13 |
| | RMS | Freq EMG | 0,11 | 0,06 | 0,16 | <0.0001 |
| **OPM** | LZ | Time | -0,48 | -0,65 | -0,32 | <0.0001 |
| | LZ | Freq OPM | 0,24 | 0,15 | 0,34 | <0.0001 |
| | LZ | RMS OPM | -0,24 | -0,37 | -0,10 | 0,001 |
| | Freq | Time | -0,61 | -0,82 | -0,40 | <0.0001 |
| | Freq | LZ OPM | 0,40 | 0,24 | 0,56 | <0.0001 |
| | Freq | RMS OPM | 0,12 | -0,06 | 0,30 | 0,18 |
| | RMS | Time | 0,81 | 0,69 | 0,94 | <0.0001 |
| | RMS | LZ OPM | -0,20 | -0,31 | -0,08 | 0,001 |
| | RMS | Freq OPM | 0,06 | -0,03 | 0,15 | 0,18 |

*Sup. Table 1. Standardized regression coefficients (β), 95% confidence intervals (CI), and FDR-corrected p-values from linear models predicting each EMG and OPM parameter during the 20% MVC condition as a function of time and the remaining parameters within the same modality.*

| Modality | Dependent | Predictor | β | CI low | CI high | $p_{FDR}$ |
|---|---|---|---|---|---|---|
| **EMG** | LZ | Time | -0,44 | -0,80 | -0,07 | 0,04 |
| | LZ | Freq EMG | 0,32 | 0,03 | 0,60 | 0,04 |
| | LZ | RMS EMG | -0,25 | -0,44 | -0,05 | 0,04 |
| | Freq | Time | -0,75 | -1,13 | -0,37 | 0,001 |
| | Freq | LZ EMG | 0,43 | 0,04 | 0,82 | 0,04 |
| | Freq | RMS EMG | 0,22 | -0,02 | 0,45 | 0,08 |
| | RMS | Time | 0,67 | 0,05 | 1,30 | 0,04 |
| | RMS | LZ EMG | -0,70 | -1,26 | -0,15 | 0,04 |
| | RMS | Freq EMG | 0,45 | -0,04 | 0,94 | 0,08 |
| **OPM** | LZ | Time | -0,03 | -0,45 | 0,39 | 0,89 |
| | LZ | Freq OPM | 0,73 | 0,38 | 1,08 | 0,001 |
| | LZ | RMS OPM | -0,24 | -0,44 | -0,04 | 0,04 |
| | Freq | Time | -0,66 | -0,91 | -0,41 | 0,0001 |
| | Freq | LZ OPM | 0,49 | 0,25 | 0,73 | 0,001 |
| | Freq | RMS OPM | 0,18 | 0,014 | 0,35 | 0,04 |
| | RMS | Time | 0,97 | 0,38 | 1,56 | 0,008 |
| | RMS | LZ OPM | -0,64 | -1,18 | -0,10 | 0,04 |
| | RMS | Freq OPM | 0,73 | 0,06 | 1,40 | 0,04 |

*Sup. Table 2. Standardized regression coefficients (β), 95% confidence intervals (CI), and FDR-corrected p-values from linear models predicting each EMG and OPM parameter during the 60% MVC condition as a function of time and the remaining parameters within the same modality.*

| Modality | Dependent | Predictor | β | CI low | CI high | $p_{FDR}$ |
|---|---|---|---|---|---|---|
| **EMG** | LZ | Protocol | 0.07 | 0.04 | 0.09 | <0.0001 |
| | LZ | Time | -0.03 | -0.07 | -0.01 | 0.009 |
| | LZ | Protocol × time | -0.08 | -0.12 | -0.05 | <0.0001 |
| | Freq | Protocol | 14.11 | 9.60 | 18.63 | <0.0001 |
| | Freq | Time | -3.36 | -7.12 | 0.04 | 0.12 |
| | Freq | Protocol × time | -14.49 | -21.30 | -7.68 | 0.0001 |
| | RMS | Protocol | -0.002 | -0.22 | 0.21 | 0.99 |
| | RMS | Time | 0.56 | 0.35 | 0.78 | <0.0001 |
| | RMS | Protocol × time | 0.24 | -0.14 | 0.61 | 0.26 |
| **OPM** | LZ | Protocol | 0.01 | -0.02 | 0.04 | 0.40 |
| | LZ | Time | -0.04 | -0.07 | -0.01 | 0.009 |
| | LZ | protocol × time | -0.05 | -0.11 | 0.001 | 0.09 |
| | Freq | Protocol | 3.27 | -0.61 | 7.15 | 0.14 |
| | Freq | Time | -4.84 | -8.11 | -1.58 | 0.008 |
| | Freq | protocol × time | -11.84 | -17.84 | -5.83 | 0.0003 |
| | RMS | Protocol | 0.15 | -0.09 | 0.38 | 0.26 |
| | RMS | Time | 0.37 | 0.13 | 0.60 | 0.005 |
| | RMS | protocol × time | -0.03 | -0.46 | 0.40 | 0.92 |

*Sup. Table 3. Standardized regression coefficients (β), 95% confidence intervals (CI), and FDR-corrected p-values from linear mixed-effects models comparing 20% and 60% MVC conditions (protocol), normalized time, and their interaction in both sEMG and OPM-MMG.*

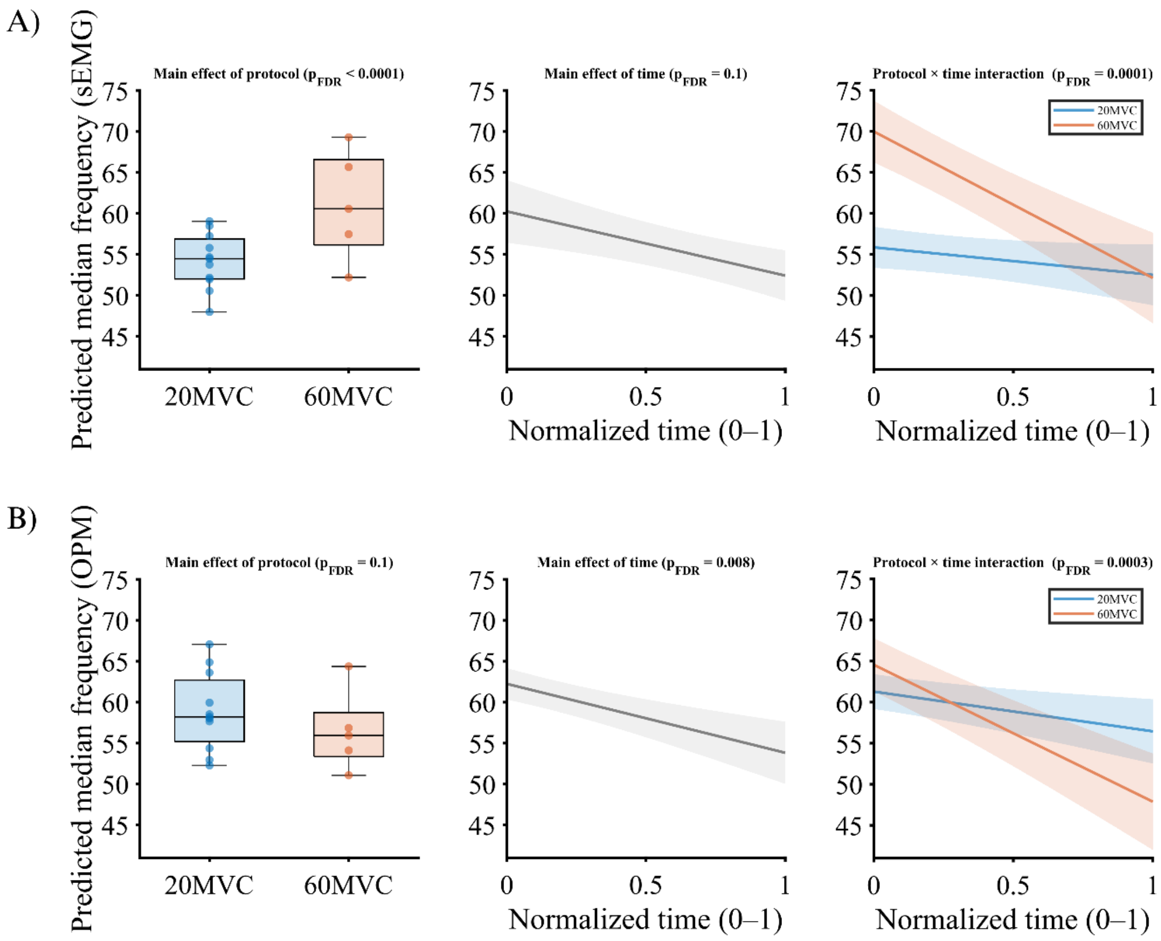


*Sup. Fig 1. Intensity- and time-dependent effects on median frequency in sEMG and OPM recordings. Predicted median frequency values derived from linear mixed-effects models are shown for surface electromyography (sEMG; A) and OPM-based magnetomyography (OPM; B). Left panels depict the main effect of protocol (20% vs. 60% MVC), middle panels show the main effect of normalized time (0-1), and right panels illustrate the protocol × time interaction. Boxplots show model-predicted values aggregated across time. Solid lines represent model-predicted trajectories, with shaded areas indicating 95% confidence intervals. False discovery rate (FDR)-corrected p-values are reported above each panel.*

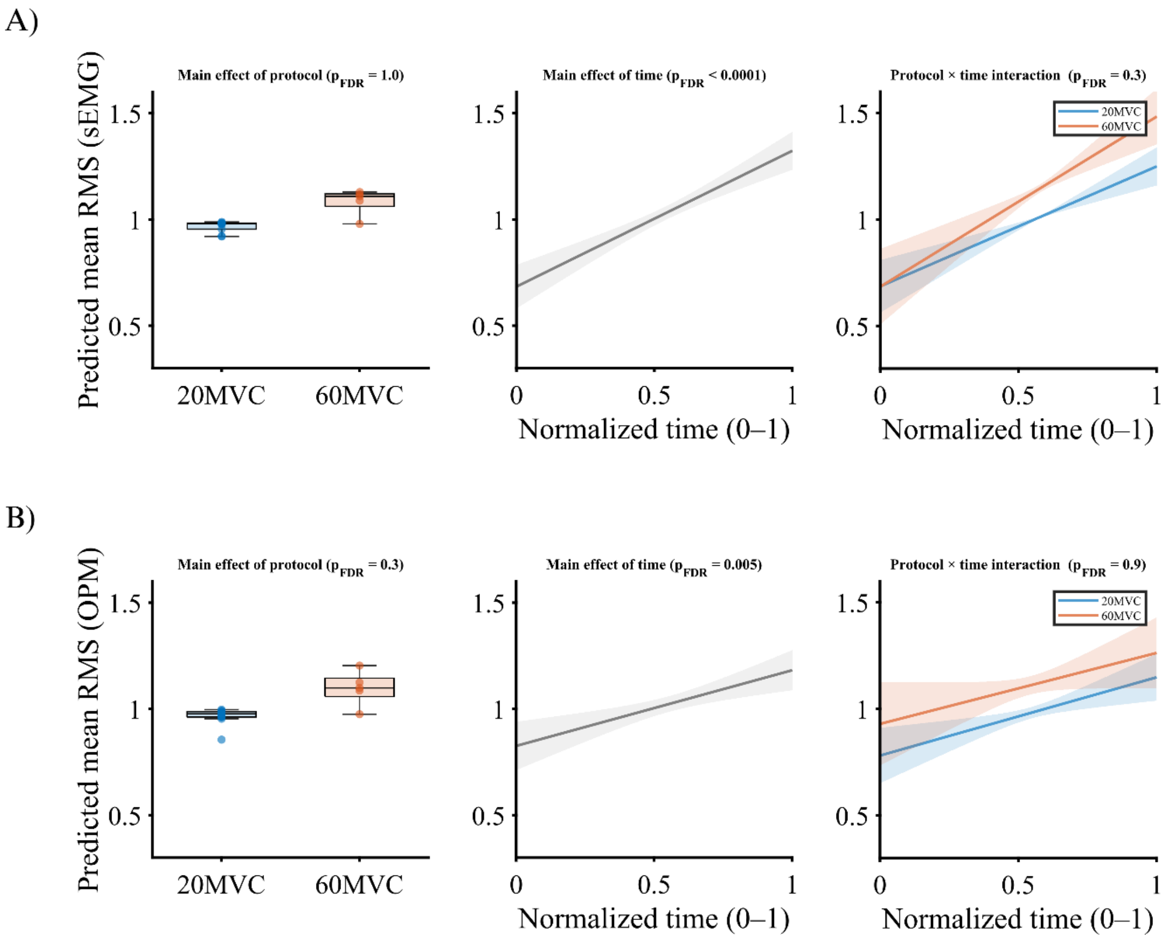


*Sup. Fig 2. Intensity- and time-dependent effects on root mean square (RMS) in sEMG and OPM recordings. Predicted RMS values derived from linear mixed-effects models are shown for surface electromyography (sEMG; A) and OPM-based magnetomyography (OPM; B). Left panels depict the main effect of protocol (20% vs. 60% MVC), middle panels show the main effect of normalized time (0-1), and right panels illustrate the protocol × time interaction. Boxplots show model-predicted values aggregated across time. Solid lines represent model-predicted trajectories, with shaded areas indicating 95% confidence intervals. False discovery rate (FDR)-corrected p-values are reported above each panel.*
